# Quadriceps-hamstrings muscle co-activation during the swing phase of walking is modulated by task constraints in healthy adults

**DOI:** 10.1101/2024.02.29.582454

**Authors:** Ellis A.M. Van Can, Han Houdijk, Tom J.W. Buurke

**Author notes:** Corresponding author Address: Antonius Deusinglaan 1, 9713 AV, Groningen, The Netherlands.

## Abstract

**Background:** Muscle co-activation, the simultaneous activation of muscles or muscle groups, is a common strategy to enhance the stability of the musculoskeletal system. However, co-activation can also be the consequence of underlying neurological impairments. To better understand and discern functional co-activation during walking, this study explored the difference in quadriceps-hamstrings co-activation during the swing phase of walking and an isolated leg-swinging movement in healthy adults.

**Methods:** Twelve healthy young adults performed walking and isolated leg-swinging at slow (0.6 m/s) and comfortable speed. Isolated leg-swinging was frequency and amplitude matched to the walk conditions. Electromyography signals from m. vastus lateralis, m. rectus femoris, m. biceps femoris, and m. semitendinosus were recorded. Pearson correlation coefficient (Pearson-CI) was calculated as a measure of rate of co-activation. Area under the curve (AUC-CI) was calculated as a measure of co-activation magnitude. Co-activation indices were calculated for both metric across the four muscle pairs and averaged into a single quadriceps-hamstrings CI for each metric.

**Results:** The results showed a higher Pearson-CI, but not AUC-CI, during walking compared to isolated leg-swinging, specifically during mid- and terminal-swing at both speeds. AUC-CI, but not Pearson-CI, was significantly higher during slow speed, compared to comfortable speed.

**Conclusion:** Quadriceps-hamstrings co-activation towards the end of the swing phase during walking reflects preparation for heel-strike, which is not present in isolated leg-swinging. Therefore, an isolated leg-swinging task could serve as a feasible method to distinguish pathological from functional muscle co-activation during walking.

## 1. Introduction

Moving around in our daily lives requires refined motor control to adapt to task and environmental constraints. Muscle co-activation, defined as the simultaneous activation of the antagonistic muscle groups, is considered to be an important motor control strategy for enhancing joint stability [1]. While an agonist muscle generates force according to the required joint torque, the co-activated antagonist generates force in the opposite direction, resulting in stiffening and thereby stabilizing the joint [2]. In this way, muscle co-activation safeguards the body against external perturbations, such as slope [3] and surface compliance [4]. During overground level walking, muscle co-activation is increased in older adults [5], individuals with musculoskeletal impairments [6,7], and individuals with neurological impairments [8,9,10]. In the latter population, in addition to being a functional strategy that enhances stability, muscle co-activation could also be a direct effect of the underlying neurological pathology [11]. In weight-bearing and thereby stability-requiring conditions, such as walking, these co-activation origins occur concurrently, making them indistinguishable.

Identifying the distinct origins of muscle co-activation may be achieved by eliminating the task constraints of walking, such as stabilization and foot placement. For instance, assessing only the swing phase in a stabilized environment allows for this assessment [12]. In theory, the swinging leg can behave like a pendulum, exchanging potential and kinetic energy and therefore requiring no muscle activation [13,14,15]. This passive energy transfer is facilitated by bi-articular muscles (e.g., m. rectus femoris, hamstring muscles) that span the hip, knee, and ankle joints [16]. However, during human walking, the swinging leg often requires active work of these muscles to control its motion as the leg swings outside its natural frequency [12]. Quadriceps and hamstrings are (co-)activated to decelerate the leg towards the end of the swing and stiffen the knee joint in preparation for weight acceptance [17,18]. Consequently, observed co-activation during the swing phase of walking could still be functional to the task constraints of walking. Although bi-articular muscles are essential for leg swinging, their simultaneous activation to control different joints complicates our interpretation.

A comprehensive understanding of the manifestation of functional co-activation in the swing phase of walking is crucial for future potential identification of pathological co-activation. This study introduces an isolated leg-swinging task that mimics the temporal (swing frequency) and spatial (swing amplitude) dynamics of the swing phase of walking, but does not require stabilization and foot placement. We hypothesize that quadriceps-hamstrings co-activation will manifest during terminal swing of the swing phase of walking as it is associated to accommodate walking-related heel strike [17,18]. Due to the absence of the task constraints in isolated leg-swinging, quadriceps hamstring co-activation will not be functional to isolated leg-swinging and hence will not manifest in able-bodied individuals. The phasing of muscle activity slightly shifts between speeds [19] and amplitude of muscle activity increases with speed [20]. This shift in phasing and change in amplitude may differ between agonist and antagonist. Therefore, we expect an effect of speed on the differences between quadriceps-hamstrings co-activation during the swing phase of walking and isolated leg-swinging.

## 2. Methods

### 2.1 Participants

Twelve healthy young adults (7 males, 5 females, age: 22.3 ±1.8 years, body height: 1.77 ±0.08 m, body weight: 72.7 ±10 kg, dominant leg: 1/11 (L/R), leg length: 0.97 ±0.04 m) volunteered to participate in this study. The inclusion criterion was age between 18 and 25 years. Exclusion criteria were (1) inability to understand the study instruction in Dutch, (2) indications of orthopedic, neurological, cardiorespiratory, and behavioral conditions that may affect walking, and (3) contra-indications for physical activity assessed by the Physical Activity Readiness Questionnaire (PARQ). The study procedures were approved by the medical ethical committee of the University Medical Center Groningen (NL83016.042.22) and in line with the Declaration of Helsinki [21]. Participants signed written informed consent before participation.

### 2.2 Experimental protocol

Participants visited the lab on a single occasion. Before the experimental trials, height, weight, and leg length were measured and participants were asked to indicate their sex and age. The study protocol consisted of four experimental conditions, two walk conditions, and two isolated leg-swinging conditions (Fig. 1) at slow walking speed (SWS), 0.6 m/s normalized to leg length [22], and comfortable walking speed (CWS).

**Fig 1.**
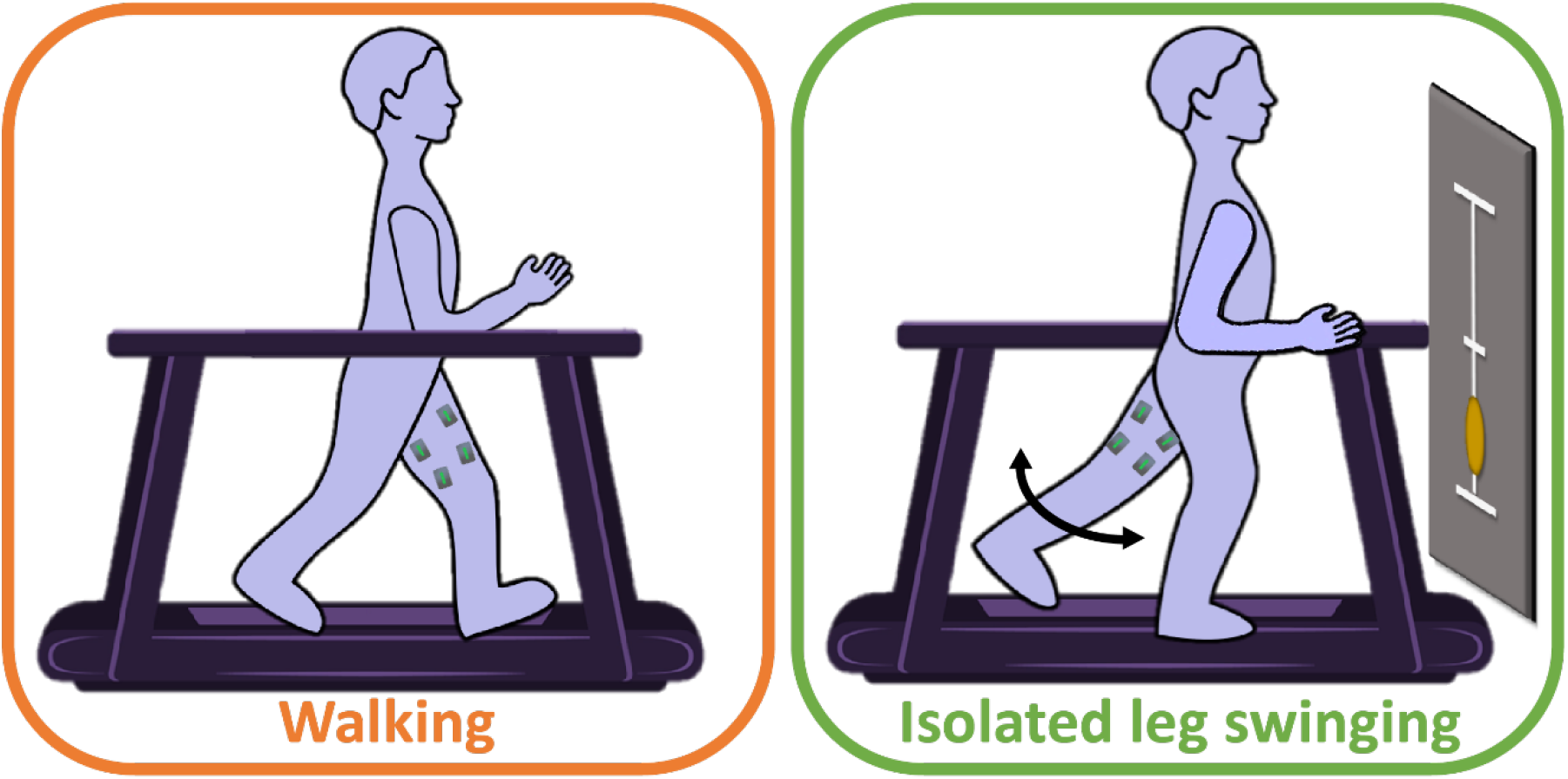
Illustration of the experimental set-up. During both conditions, electromyography signals of quadriceps and hamstrings, kinematic and kinetic data were recorded. Motion capture cameras and passive markers on trochanter major and lateral malleolus are not included in the figure.

Participants walked on the instrumented treadmill in a six-minute familiarization trial during which their CWS was established. Participants were asked to indicate their CWS while the experimenter increased the belt speed gradually. The belt speed was then set to 0.3 m/s above the indicated CWS while the experimenter gradually decreased the belt speed, participants were again asked to indicate their CWS. The CWS used in the experiment was the highest indicated speed in the two trials.

In the walking conditions participants were asked to walk for four minutes on the treadmill. Participants were not allowed to touch the handrails and were fitted with a safety harness to prevent falls, without providing support. During the last minute of the walking condition, the mean swing frequency was calculated from the mean time from toe-off to heel-strike, and the corresponding mean swing distance was calculated. In the subsequent isolated leg-swinging conditions, participants had to swing their dominant leg at the calculated swing frequency indicated by a metronome. Visual feedback on the swing distance was provided by a projection on the screen in front of the treadmill.

The isolated leg-swinging condition consisted of four 50-second swing periods with 10-second rest periods in between to prevent fatigue. Participants stood in an upright stance position and were allowed to lean slightly to the side for clearance. Participants were instructed to swing their leg without touching the surface while keeping the metronome pace. Visual feedback on the swing amplitude was provided on a screen in front of the participants (Fig 1). They were allowed to rest their arm, contralateral to the swinging leg, on the handrail that was adjusted to their elbow height. Participants wore a safety harness during all conditions and received a minimum rest period of three minutes in between experimental conditions. The sequence of the two paired experimental conditions (SWS_walking_ – SWS_isolated leg-swinging_ and CWS_walking_ – CWS_isolated leg-swinging_) was randomized between participants. Fig 2 shows one possible sequence of conditions.

**Fig 2.**
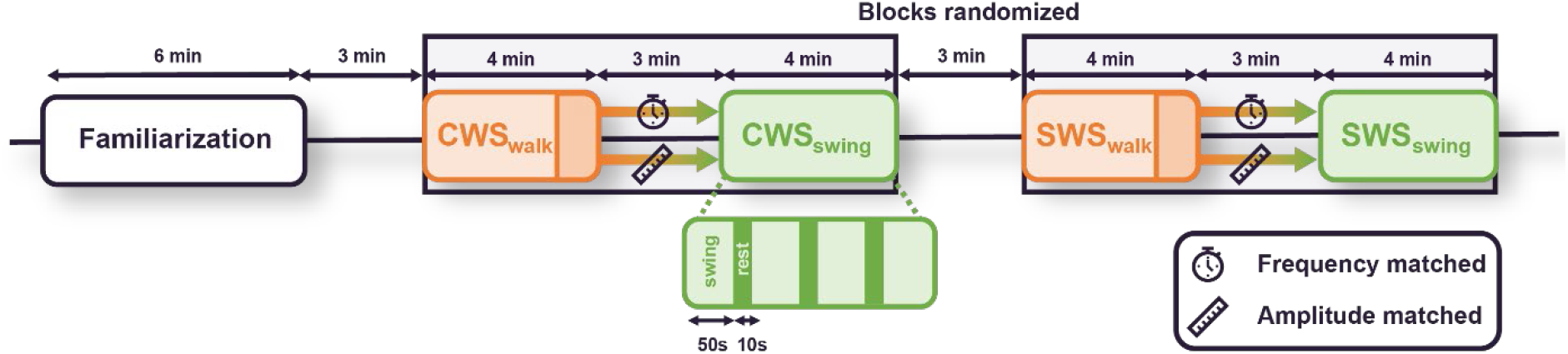
Timeline of experimental conditions. Orange blocks represent the walking conditions (‘walk’), and darker orange blocks represent the last minute in which to the frequency and amplitude of leg swing were calculated. Green blocks represent the isolated leg-swinging conditions (‘swing’). Both swing conditions consisted of 4 times 50 seconds leg-swinging periods (light green in example) followed by 10 seconds rest (dark green). Blocks of walking and isolated leg swinging at the two speeds were randomized. In between each condition was a 3-minute rest period. CWS: Comfortable walking speed. SWS: Slow walking speed.

### 2.3 Instrumentation & data collection

Electromyography (EMG) data were recorded using bipolar surface electrodes (Delsys 16-channel sEMG system, Natick, MA, USA, electrode size: 27×37×13 mm inter-electrode distance: 10 mm) at a sampling frequency of 2148 Hz. Electrodes were placed on four lower extremity muscles of the dominant leg (quadriceps: m. rectus femoris (RF), m. vastus lateralis (VL ); hamstrings: m. biceps femoris (BF)). After the skin surface was shaved and cleaned with alcohol, the electrodes were placed according to the SENIAM conventions [23]. A three-dimensional motion capture system (10 cameras; Vicon Motion Systems Ltd, Yarnton, UK) recorded the trajectory of the lateral malleolus and trochanter major of the dominant leg at a sampling frequency of 100 Hz. Three-dimensional ground reaction force (GRF, N), Center of pressure (COP, m), and three-dimensional moment (Nm) were measured with two force plates embedded in the treadmill (Motek Medical, Amsterdam, the Netherlands) at a sampling frequency of 1000 Hz. EMG, kinematic, and kinetic data were time-synchronized through a software trigger.

### 2.4 Data analysis

All data and statistical analyses were performed in MATLAB (version 2023b; The MathWorks Inc. Natick, MA, USA). The raw EMG signals were bandpass filtered with a 2^nd^ order Butterworth filter (20-500 Hz [23] and full wave rectified and low-pass filtered (2^nd^ order Butterworth filter, 20 Hz [23]). The signals were visually checked for artifacts. EMG amplitude normalization was done with respect to peak amplitude overall conditions [24]. After normalization, the first and last five seconds of each signal were removed, to eliminate muscle activation due to initiation and termination of walking and swinging.

The COP data were high-pass filtered (5 Hz, 2^nd^ order Butterworth filter), detrended and low-pass filtered (10 Hz, 2^nd^ order Butterworth filter)[25]. The first and last five seconds of the COP signal were removed. The peaks in the anterior-posterior COP signal were used to detect foot contact events. The swing phase in the walking condition was defined as toe-off to heel-strike of the dominant leg. For isolated leg-swinging, the swing phase was defined as the maximal posterior position of the ankle to its maximal anterior position. EMG data were resampled from toe-off to ipsilateral heel-strike for walking and from posterior to anterior for isolated leg-swinging, respectively, on a 100-point time base. The four resampled EMG signals were averaged to one single swing EMG signal for each muscle. The first and last five seconds were removed for each of the four swing instances during walking and isolated leg-swinging.

Co-activation index (CI) was calculated for the quadriceps-hamstrings pairs (QD-HS; RF-BF, RF-ST, VL-BF and VL-ST) using Pearson correlation coefficient to reflect the direction and strength of the linear relationship of two muscles (Pearson-CI, Eq. 1)[26,27], in this study we refer to Pearson-CI as the rate of co-activation. Positive Pearson-CI indicated co-activation, while negative Pearson-CI indicated no co-activation, i.e. the two muscles oppose each other in activation (Fig. 3). In addition, area under the curve (AUC) was included as a measure of the absolute quantity of the co-activation. We calculated AUC as the cumulative shared area at each time instance as a percentage of the maximal area of that specific time instance (adapted from [28]). The four quadriceps-hamstrings Pearson-CIs and AUC-CIs were averaged to one QD-HS Pearson-CI and one QD-HS AUC-CI for each time interval of interest (full, initial, mid, terminal swing) in each condition.

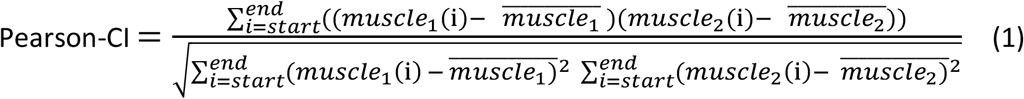

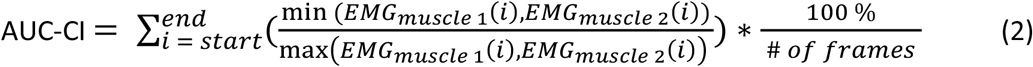

**Fig. 3.**
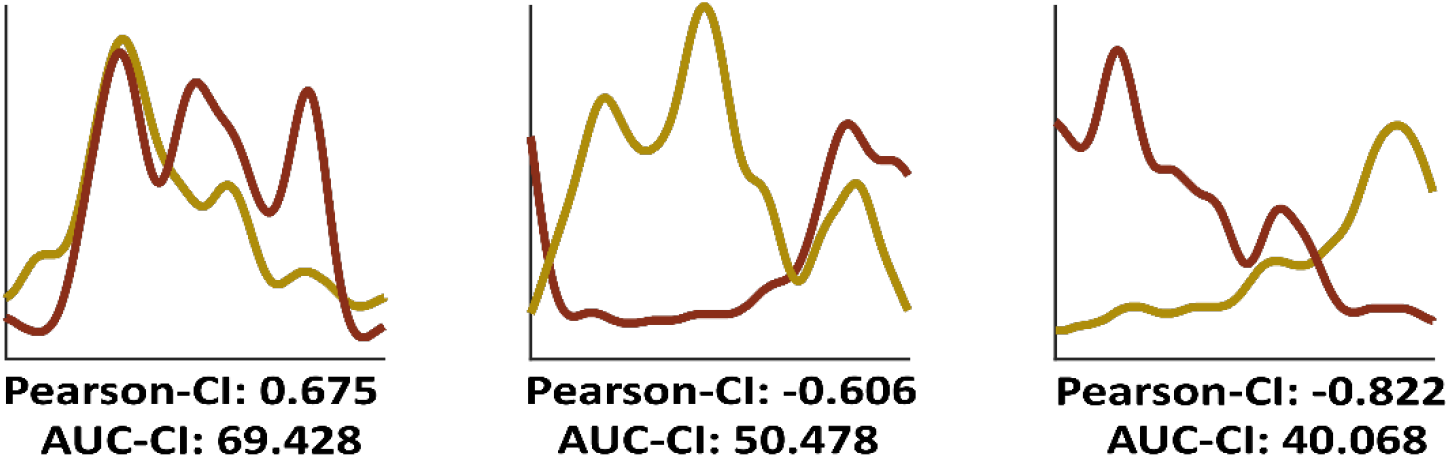
Example of Pearson and area under the curve (AUC) co-activation indices (CI). Note that muscle activation and indices in the figure serve as examples and do not portray actual gait or leg-swinging data.

### 2.5 Statistical analysis

To assess whether swing time during isolated leg-swinging matched swing time during walking we performed two paired t-tests to compare swing time between walking and isolated leg-swinging for SWS and CWS separately.

Differences in co-activation of QD-HS between the walking and isolated leg-swinging conditions were tested using two (Pearson-CI, AUC-CI) two-way (2*2) repeated measures ANOVAs with condition and speed as within-subjects factors. In case of violation of sphericity, Greenhouse Geiser corrected p-values were interpreted. Significant interaction effects were evaluated with Bonferroni post hoc corrections. Eta-squared (η^2^) was reported and interpreted as 0.01 small effect size, 0.06 medium effect size 0.14 large effect size [29].

To investigate in which part of the swing phase differences in co-activation between walking and isolated leg-swinging occurred, we first calculated a moving average of Pearson-CI and AUC-CI, using time bins each representing 25% of the swing phase. Then, four Statistical Parametric Mapping (SPM) paired t-tests (for each outcome (Pearson-CI and AUC-CI) and each speed) were performed on the moving average signals. SPM is a technique that allows for the statistical analysis of temporal signals [30]. Statistical significance was set at p < 0.05 for all tests.

## 3. Results

The mean comfortable walking speed was 1.21 ±0.17 m/s and slow walking speed was fixed at 0.6 m/s normalized to leg length [22]. SWS walking swing time (0.46 ±0.09 s) was not significantly different from SWS isolated leg-swinging swing time (0.52 ±0.06 s, t(1,11)=-1.821, p=0.096). Furthermore, CWS walking swing time (0.39 ±0.02 s) was not significantly different from CWS isolated leg-swinging swing time (0.41 ±0.04 s, t(1,11)=-1.556, p=0.148).

### 3.1 Quadriceps-hamstrings co-activation during the full swing phase

Quadriceps and hamstrings muscle activation varied depending on the condition and speed (Fig. 4). Pearson-CI was significantly higher in the swing phase of walking (0.477 ±0.253) compared to isolated leg-swinging (-0.310 ±0.341; F(1,11)=62.131, p<0.001, η^2^=0.642; Fig. 5), but AUC-CI was not different (Swing phase walking: 51.010 ±9.867, isolated leg-swinging: 46.545 ±13.668; F(1,11)=6.235, p=0.030, η^2^=0.035). AUC-CI was significantly higher during SWS (54.740 ± 12.164) compared to CWS (42.814 ± 8.51; F(1,11)=31.296, p<0.001, η^2^=0.252), but speed did not have a significant effect on Pearson-CI (SWS: 0.137 ±0.523, CWS: 0.029 ±0.472; F(1,11)=2.450, p=0.146, η^2^=0.012). Finally, no significant condition*speed interactions were found for Pearson-CI (F(1,11)=0.868, p=0.371, η^2^=0.004) or AUC-CI (F(1,11)=2.215, p=0.164, η^2^=0.016).

**Fig. 4.**
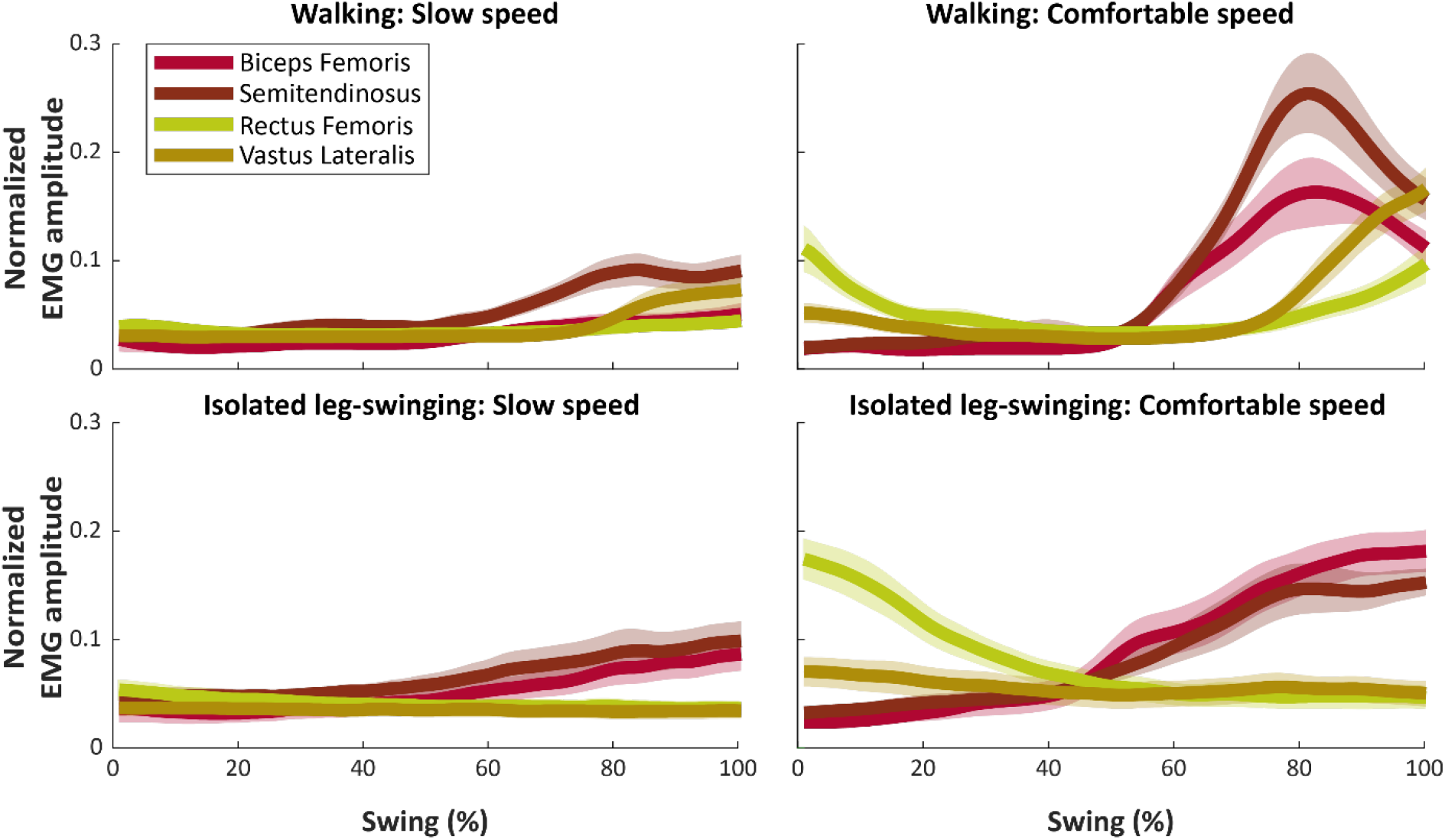
Time normalized group-averaged muscle activity. Shaded areas around the mean represent the standard error of the mean.

**Fig. 5.**
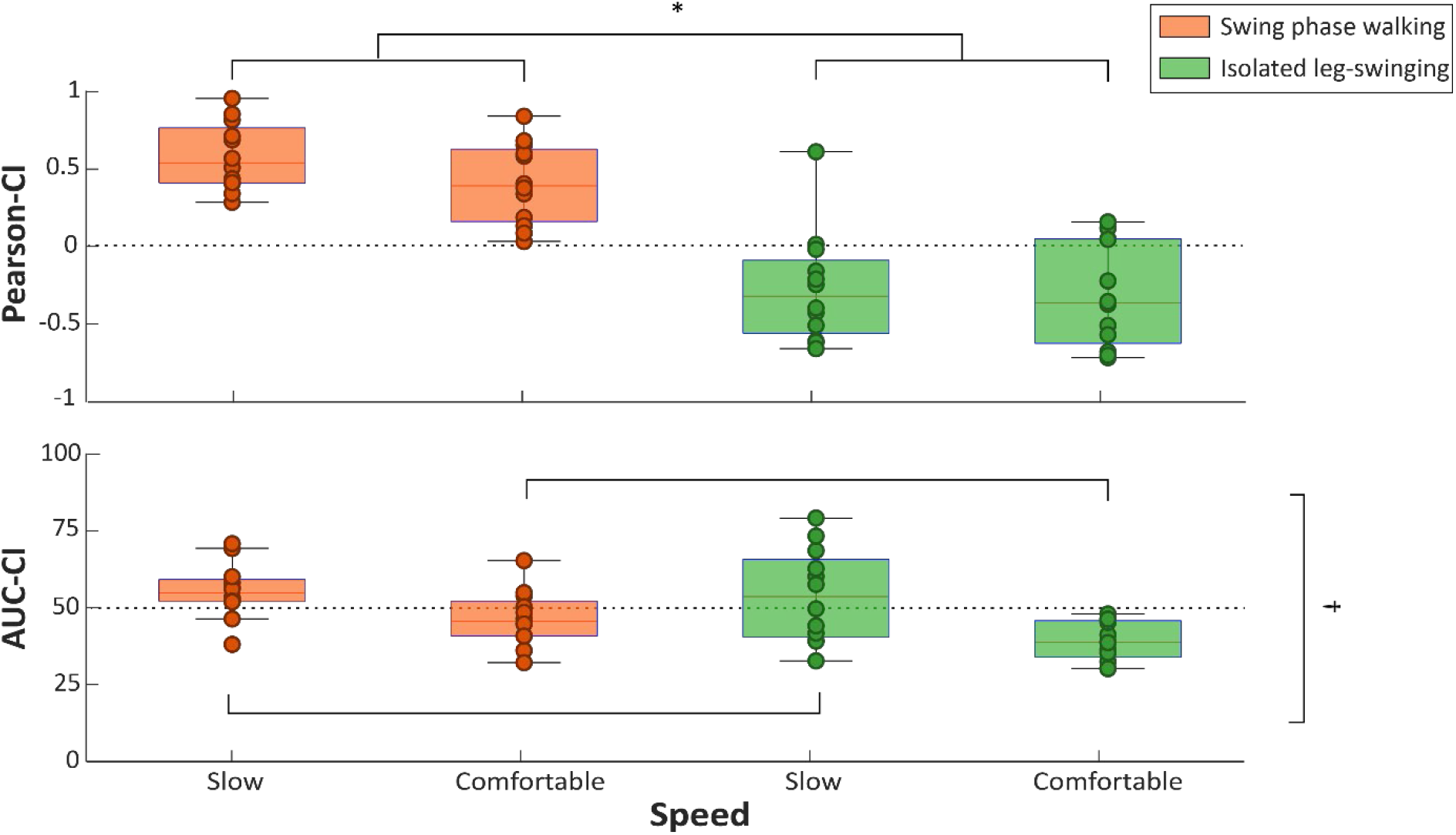
Pearson and area under the curve AUC co-activation indices (CI) for slow walking speed (SWS) and comfortable walking speed (CWS). * Indicates significant main effect for condition, † indicates significant main effect for speed.

### 3.2 Differences in quadriceps-hamstrings co-activation throughout subphases of swinging

The SPM paired t-tests (Fig. 6) showed significantly higher Pearson-CI in SWS walking than SWS isolated leg-swinging for windows 69-81 (t(1,11)=3.380, p<0.001). Furthermore, Pearson-CI was significantly higher in CWS walking than CWS isolated leg-swinging for windows 52-57, windows 59-60, and windows 55-74 (t(1,11)=3.454, p=0.015, p=0.048, p<0.001 respectively). Finally, AUC-CI was significantly higher during CWS walking than CWS isolated leg-swinging for windows 85-87 (t(1,11)=2.974, p=0.048).

**Fig. 6.**
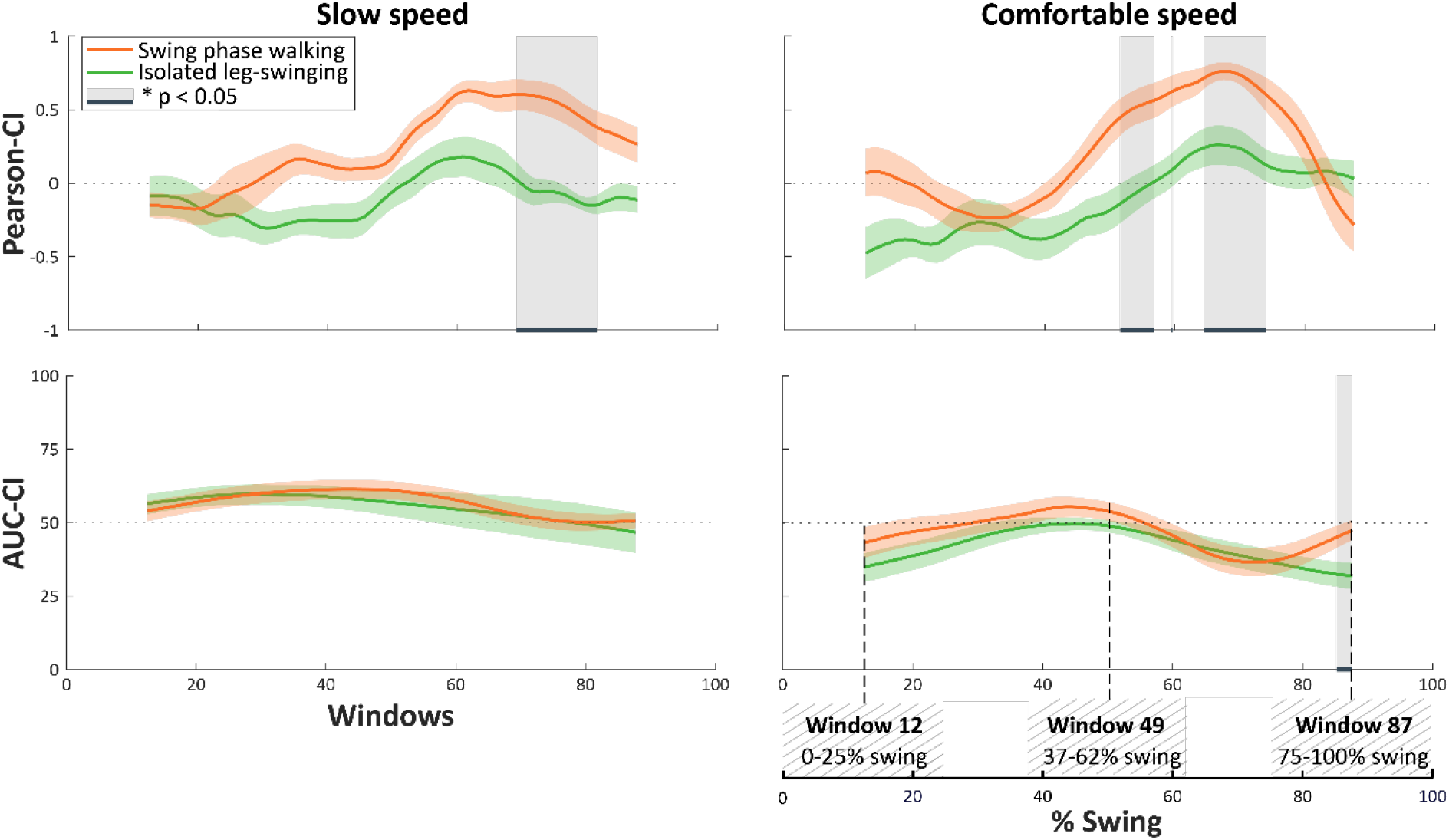
Statistical parametric mapping (SPM) results showing group averaged Pearson and area under the curve (AUC) co-activation indices (CI) for each window representing 25% of both swing conditions. Shaded areas around the mean represent the standard error of the mean. The lower-right panel indicates how three windows represent percentages of the original swing phase.

## 4. Discussion

This study aimed to investigate the differences in quadriceps-hamstrings co-activation between the swing phase of walking and isolated leg-swinging. The results showed higher rate of co-activation during walking compared to isolated leg-swinging. The differences in QD-HS co-activation rate were apparent for SWS in windows representing 57-94% of the swing phase and for CWS in windows representing 40-70%, 47-73%, and 43-87% of the swing phase. The magnitude of co-activation was different in windows representing 57-94% of the swing phase. The results showed no effect of speed on the rate of co-activation but magnitude of co-activation was higher at slow speed, compared to comfortable speed. These results indicate that the rate of QD-HS co-activation is different during mid and terminal part of the swing phase and the magnitude of co-activation is different towards and during terminal swing.

As hypothesized, QD-HS co-activation was observed during walking but not during isolated leg-swinging. During both the swing phase of walking and isolated leg-swinging, activation of quadriceps muscles propels the swinging leg forward. This forward motion is decelerated by activation of the hamstrings [18]. Towards the terminal swing of walking, quadriceps-hamstrings co-activation prepares weight acceptance of the swinging leg by stabilizing the knee joint [18,31]. Although able-bodied individuals can selectively control their hamstrings and quadriceps muscles, our results show that this cannot be inferred from muscle activation during the swing phase of walking alone. The selective control of quadriceps and hamstring muscles could only be observed during isolated leg swinging when the stabilization and foot placement constraints from walking were removed.

The main effect of speed on the magnitude of co-activation indicates that there was more quadriceps-hamstrings co-activation at slow speeds compared to comfortable speeds. This could be explained by two different phenomena. First, postural stability demands are higher at lower speeds, which could increase muscle co-activation to accommodate these demands. Second, low activation of all muscles at slower speeds might inflate the AUC-CI. AUC-CI shows a co-activation of approximately 50% throughout both conditions despite lower muscle activations at lower speeds (Fig 4, Fig 5). AUC-CI therefore measure expresses the quantity of muscle activation relative to maximal amplitude at the same time instance. Therefore the ongoing diminished muscle activation during SWS conditions compared to CWS conditions, is reflected in relatively high co-activation which resulted in a significant difference between the two conditions.

In this study, the measure of magnitude (AUC-CI) seems to be less sensible to detect differences between the two conditions than the measure of rate (Pearson-CI). The disparities between these two measures demonstrate that whether muscles are classified to be co-activated or not is highly dependent on the methodological decision for the co-activation metric. This raises questions on how muscle co-activation could be defined. Pearson-CI indicates whether the activation of a muscle is associated with the change in the activation of another muscle. This association is either in the same direction (positive CI: co-activation) or in the opposite direction (negative CI: no co-activation [32]). This metric is robust against any amplitude normalization method but unlike other methods [10,28,33,34] does not take the relative timing and magnitude of the co-activation into account. In addition, one could argue that two muscles co-activated in the same direction but with only a small magnitude contribute to a relatively small part of the total force production [35]. To indicate the relative magnitude of the co-activation, we expressed co-activation in an additional AUC-CI. However, as mentioned before, AUC-CI may become inflated in instances in which both muscles are of relatively low amplitude. Consensus on the the use of co-activation metrics remains elusive, primarily due to the broad definition of co-activation. The results of this study further emphasize a clear definition of co-activation and subsequently cautious selection of suitable metrics in future studies.

A methodological consideration was the use of averaged quadriceps co-activation from four distinct individual QD-HS muscle pairs. m. biceps femoris, m. semitendinosus, and m. rectus femoris are bi-articular muscles and act on the knee and hip joint, while the m. vastus lateralis only spans the knee joint. This approach could potentially oversimplify the complexities of thigh muscle co-activation during swing. However, this consideration increases this study’s statistical power as the focus was the difference in general quadriceps-hamstrings muscle co-activation during walking and isolated leg-swinging.

In future work, our isolated leg-swinging task could serve as a potentially tool in clinical populations to reveal co-activation as the result of impairments. Currently, it is unclear to what extent increased muscle co-activation in clinical populations could be attributed to underlying deficits, limiting treatment selection. For instance, in individuals post-stroke, stiff knee gait (i.e. hip circumduction), a frequently observed gait deviation during the swing phase, could be the direct effect of impaired motor control of knee extensors [36], but could also be a secondary compensation for reduced push-off power of ankle muscles [37]. In this population observed co-activation during the isolated leg-swinging task would indicate that this co-activation is inherent to their impairments rather than a compensatory strategy.

It can be concluded that quadriceps-hamstrings co-activation towards the end of the swing phase of walking reflects functional co-activation related to simultaneous deceleration of the leg and joint-stiffening in preparation for heel-strike during walking. The isolated leg-swinging task presented in this study allows for the differentiation of pathological muscle co-activation from task-related functional muscle co-activation during walking in the future. It should be taken into account that the degree of co-activation is highly influenced by the selected co-activation metric, which emphasizes the need for clear and cautious considerations on the definition, selection and interpretation of co-activation metrics.

## Acknowledgements

The authors wish to thank Anniek Heerschop, Emyl Smid, Dirk van der Meer, Wim Kaan, Daniel Dias dos Santos, David Hozeman, Isa Kreuwel and Marije Bouius for their help.

